# scDNA: Single Cell DNA analysis software toolkit for subclonality discovery and assessment

**DOI:** 10.64898/2025.12.19.694255

**Authors:** Michael Bowman, Shreeya Gounder, Varsha Singh, Olga Shestova, Troy Robinson, Amy Zhang, Anushka Gandhi, Roopsha Bandopadhyay, Sheng F. Cai, Ross L. Levine, Saar I. Gill, Linde A. Miles, Robert L. Bowman

## Abstract

Advances in single cell multi-omics technologies have allowed for investigation into genotype-immunophenotype relationships and dynamic clonal changes in cancer patient samples. These technologies provide rich insights into genetic profiling, drivers of disease progression, and subclonal dynamics. Increased adoption and throughput of these technologies, necessitate accessible computational tools for processing and analysis. Toward developing easy to use computational tools, we introduce the <u>S</u>ingle <u>C</u>ell <u>DNA</u> (scDNA) package that allows for rapid analysis of single-cell molecular profiling. Our platform aims to provide a diagnostic summary tool for sample quality control, representative subclonal architecture within a sample, demultiplexing functionality for multi-sample processing, copy number variation, and mutation trajectory analysis to identify order of subclonal mutation acquisition. We showcase a series of vignettes on hematopoiesis datasets for each aim that reflects recently deployed uses of scDNA. Additionally, we show scDNA provides a modular, user-friendly framework that readily feeds into other standard software pipelines.

## INTRODUCTION

Advances in single cell technologies have allowed for insights into subclonal properties not readily available by bulk sequencing. Easy to use bioinformatic tools are needed for rapid analysis of patient samples to investigate genotype-immunophenotype relationships and dynamic clonal changes^1^. Single cell RNA-sequencing analysis has been accelerated by accessible computational frameworks including the R-based Seurat ^2^ and python-based scanpy^3^. However, there are limited tools for single cell DNA-sequencing^4–6^ analysis that integrates into these software ecosystems. We and others have applied single cell DNA sequencing approaches to investigate clonal architecture mutation order in leukemic evolution^7–11^, genome editing^12^, measurable residual disease testing ^13, 14^, and longitudinal response to therapy^8, 15, 16^. Specifically, we have deployed computational methods for inference of clonal architecture, size, and diversity ^7, 15^. However, there are still significant gaps in our ability to assess cell quality, technical allele dropout, cell type-specific mutation enrichment, and copy number analysis.

To address these limitations, we aimed to develop a software package that comprehensively allows 1) interrogation of genetic variation at single cell resolution, 2) determination of clone number, size, and diversity, 3) identification of patterns of co-mutation, mutual exclusivity, abundance in dominant vs minor clones, and 4) determination of whether genetic alterations, alone or in combination, are associated with alterations in cellular immunophenotype.

Toward this effort, we introduce the R-package <u>S</u>ingle <u>C</u>ell <u>DNA</u> (scDNA) which allows for rapid analysis of single cell DNA-sequencing data. Our package contains utilities to determine overall quality of samples and in-depth analysis of clonal landscapes (**Figure 1**). Our pipeline automatically annotates variants from standard mutation analysis output files (VCF or Tapestri H5) and allows users to determine select variants of interest for targeted clonal analysis. In addition to mutation annotation, quality control is performed for both Variant Allele Frequency (VAF) and genotyping frequency across cells. We also provide computational tools for de novo demultiplexing of pooled samples without the need for externally validated sample-specific features. After sample assignment, clonal architecture and abundances are identified. scDNA then performs downstream analysis based on user needs including cell immunophenotypic labelling, statistical analysis, copy number variation, and clonal trajectories.

**Figure 1.**
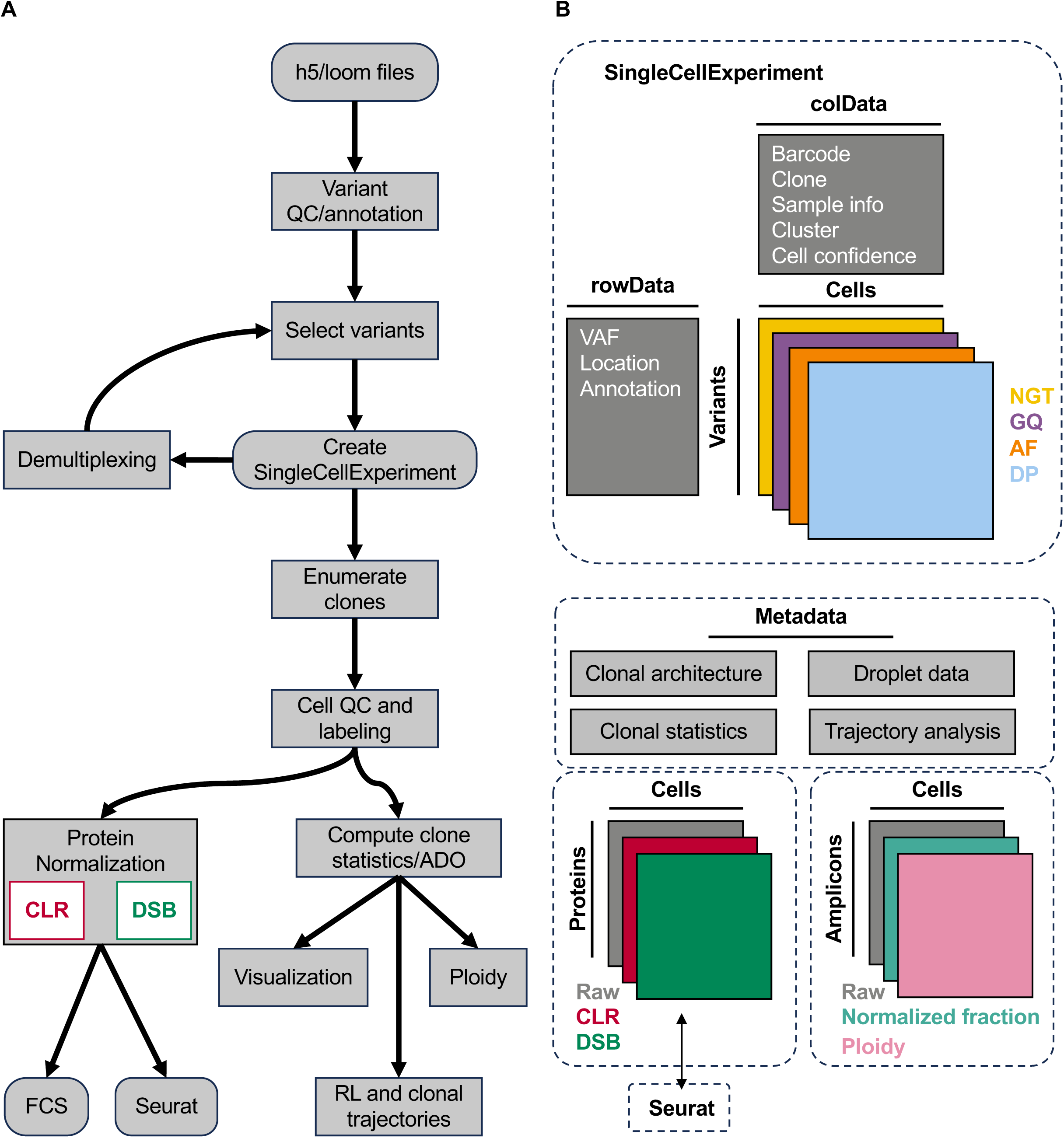
The overall pipeline and storage of data using scDNA package. **A)** contains the pipeline for standard analysis. Users have abilities to select the variants of importance for their analysis. Afterwards data is converted into a SingleCellExperiment object. Downstream analysis includes clonal architecture and find cell quality whereafter analysis can be split to potential avenues including downstream immunophenotyping, trajectory analysis, and copy number variations. **B)** Architecture of SingleCellExperiment object and data layers. Cell ID (barcode) is on the columns, and variant information is on the rows. We contain layers of matrices foe each cell’s read depth (DP), Genotype calls (NGT), genotyping quality (GQ), and Allele Frequency (AF). We also contain information regarding clone specific results. Lastly, we contain alternative experiments to keep track of protein data and copy number variation for downstream analysis.

We employ the SingleCellExperiment^17^ container to manage the different layers of data. The advantages of this framework allow for 1) straightforward mapping between a cell barcode and mutational profile, and 2) easy integration with other downstream pipelines including Seurat. The multiple assay format allows for easy extension to new multi-omic datasets including DNA+Protein^18, 19^, DNA+DNAmethylation^20, 21^, DNA+CNV^6^, and DNA+RNA^22, 23^ (**Figure 1B**). The flexible nature of the metadata slot allows for storage of descriptive summary data including clonal architecture, statistical analysis of clone size and diversity, and mutational trajectory analysis. This framework adds flexibility for future developments in bioinformatic packages, as well as future developments in the scDNA package.

## RESULTS

### Overview

Several of the standard functionalities of the scDNA package have been previously deployed in our prior work including variant identification, clonal abundance assessment, clonal diversity, clonograph visualization, and integration of DNA+Protein analysis ^7, 13, 15^, 24. Analysis begins with input from formatted h5 files (Mission Bio Tapestri Pipeline) or standardized multiVCF files. Downstream output includes determination of VAF, genotyping quality, clonal architecture and statistics. Specifically, this package contributes the following: 1) diagnostic summary tools for sample and cell quality assessment, 2) clonal architecture representation for subclonality within a sample, 3) computational demultiplexing strategies, 4) locus-specific, allele drop out aware, clonal trajectory analysis, and 5) integration with existing single cell software. We demonstrate the utility and capabilities of the package through a series of vignettes on new and previously published datasets.

### Droplet and cell quality control

Our pipeline addresses quality control at multiple stages, particularly stringent filtering on variant identification. Critical filters include read depth, allele frequency, and genotyping quality. Cells that pass all internal quality control checks are labeled as “Complete” while cells that fail are labeled as “Other.” This latter group represents cells where successful genotype calls were made^25^, but failed stringent internal filters for at least 1 variant of interest. These stringent genotyping controls ensure the identified clones comprise the highest quality cells for downstream analysis. Lastly, we have additional pre-built QC measures and clonal statistics that are accessible in **Supplemental Table 1** with the description of their capabilities and primary utility. Once cells are selected based on quality, genotype-immunophenotype specific analysis is performed.

In addition to variant QC, we propose new methods to quantify cell quality from droplet information. Tapestri pipeline typically calls cells based on DNA amount and overall amplicon coverage^19^. We introduce using protein content into our strategy for cell confidence calling. Our rationale for including protein content is that true cells should contain high levels of DNA and protein compared to ambient background droplets. We assessed total DNA and protein levels on a PBMC DNA+Protein sample stained with the Total Seq-D antibody panel (**Figure 2A**). Initial cell type calls demonstrate a clear separation between empty droplets (red) and true cells (green), with true cells possessing greater DNA and protein content, matching expectations. However, there remain empty/partial droplets which appear to have degraded DNA or incomplete panel coverage, despite ample protein content. These droplets can overlap with high quality cells with complete information and are distinct from the clearly empty droplets which have diminished coverage for both DNA and protein. This partial droplet group is critical to identify for downstream protein analysis as they likely represent poor-quality cells and not background ambient noise in empty droplets, potentially complicating cell type and mutation zygosity calls. To further refine cell quality categorization, we employ an outlier score-based cell confidence algorithm on high-quality cells using both DNA and protein information (see Methods). We assessed the outlier scores for different aggregated information of each cell including the total protein content and the number of uniquely identified cell surface markers found (**Figure 2B**). This approach identifies that poor-quality cells can arise because of poor DNA coverage or high variance sampling of protein coverage. While low protein coverage may indicate inadequate amplification of the protein library, high protein coverage might be a result of non-specific antibody accumulation in dead cells, or multi-cellular aggregates (doublets). Both cases of extremes are identified as low confidence as they complicate cell type identification in downstream clustering.

**Figure 2.**
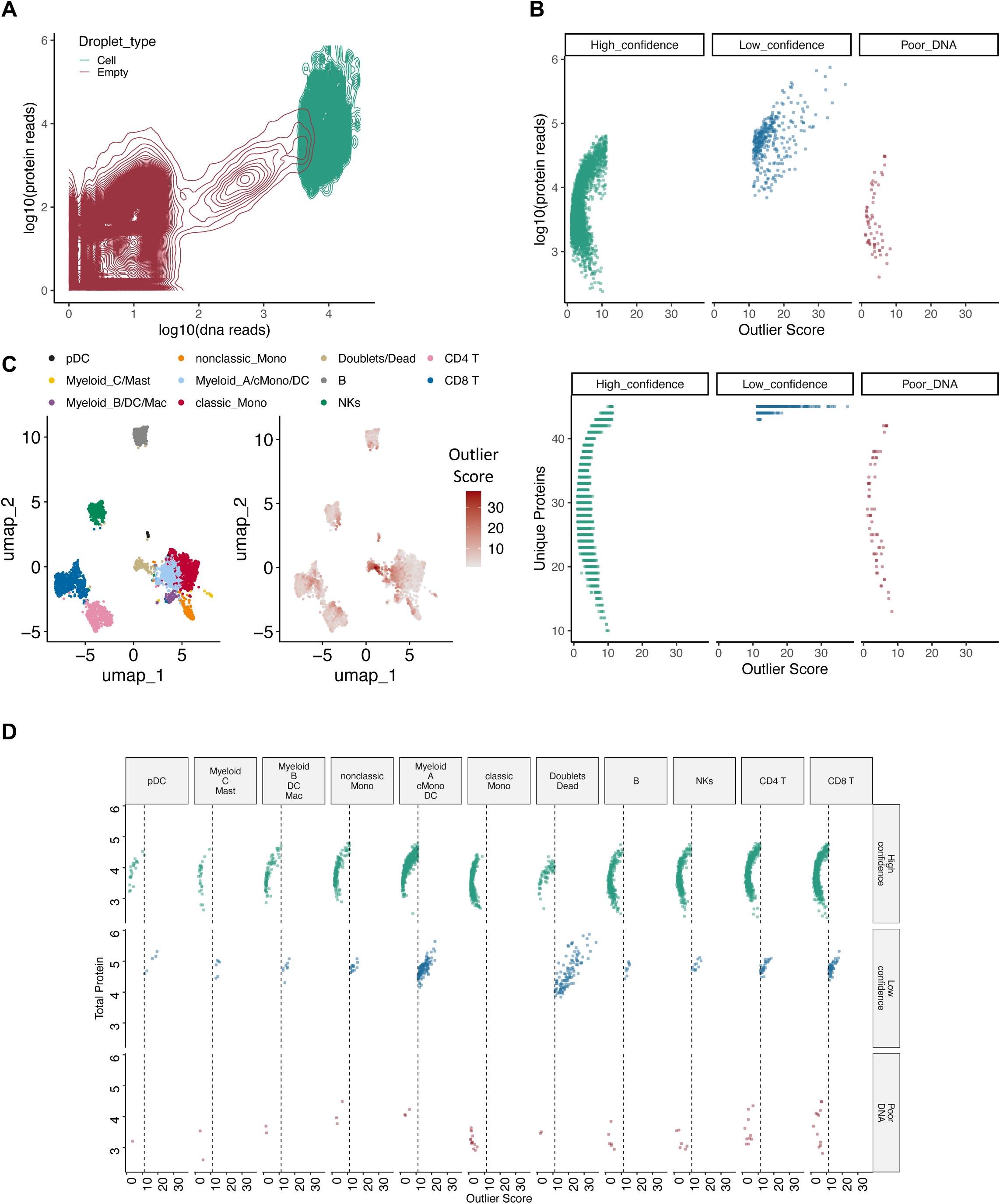
Integrating DNA and protein metrics for cellular quality assessment. **A)** Contour plot of droplets distribution of total DNA and protein reads with empty droplets (red) and true cell calls (green). **B)** Dotplot of external outlier score metric (x-axis) and total protein content (y-axis, top) and unique markers seen in a cell (y-axis, bottom). Dots depict cells originally identified by Tapsetri pipeline. The outlier score is higher for droplets with exceedingly high and low protein content to give the characteristic boomerang shape. Low confidence cells have nearly all markers present, a hallmark of associated with permeated membranes of cells or doublets. Poor DNA is based on entropy metrics from droplets that contain the 90% coverage of amplicons. **C)** UMAP of Human peripheral blood sample with manually curated cell identities and an outlier score overlay. High values concentrate on the doublet population which tend to have higher levels of protein and DNA content. **D)** The cell type breakdown for the different classes of cell quality scores. Doublets have the most low confidence cells while most cells are in high confidence category.

We next sought to determine if our cell confidence calls were cell type dependent. We clustered cells based on Protein data, identified cell types by manual curation, and overlaid cell confidence scores (**Figure 2C**). We assessed the distribution of poor DNA quality, low confidence and high confidence across these cell populations. We identified one cluster with high overlap with low confidence cells and increased total protein, indicative of dead cells (**Figure 2D**). We did not observe cell type skewing toward low confidence cells that might arise because of the antibody panel design. For instance, low-confidence cells were not restricted to lymphoid cells due to an imbalance in lymphoid-myeloid marker proteins on the panel. Conversely, greater myeloid marker representation on the panel did not artificially identify low-confidence cells with high protein expression.

### Cell type enrichment for mutations and clones

Our pipeline can determine mutant allele frequency and clone distribution across entire samples and within specific cell subsets. To demonstrate this utility, we analyzed scDNA and protein expression from an AML patient sample harboring mutations in *TET2, NPM1* and *NRAS,* identifying broad classes of leukemic stem cell (blast), monocytic and lymphoid populations^15^ (**Figure 3A**). When approaching the entire sample, we take a “variant-first” approach where mutations of interest are first identified and enumerated into clones. These clones are then assigned as metadata, and their abundance can be assessed across cell type designated by the protein data. These clones can then be overlayed on the identified cell types to visualize trends in clone-cell overlap, including an abundance of non-mutant, wildtype (WT) clones in lymphoid (**Figure 3A).** We additionally, observe the *TET2-NRAS* clone appears more abundant in the monocytic populations, while the *TET2*-*NPM1* clone is expectedly in the hematopoietic stem cell progenitor compartments. As a compliment to identifying overlapping trends of clones and cell clusters, we can also perform differential marker expression between clones. We observed higher enrichment for CD14 and CD11b in *TET2-NRAS* compared to *TET2-NPM1,* indicating increased myeloid differentiation *in NRAS*-mutant cells as we and others previously described ^7^ **(Figure 3B**).

**Figure 3.**
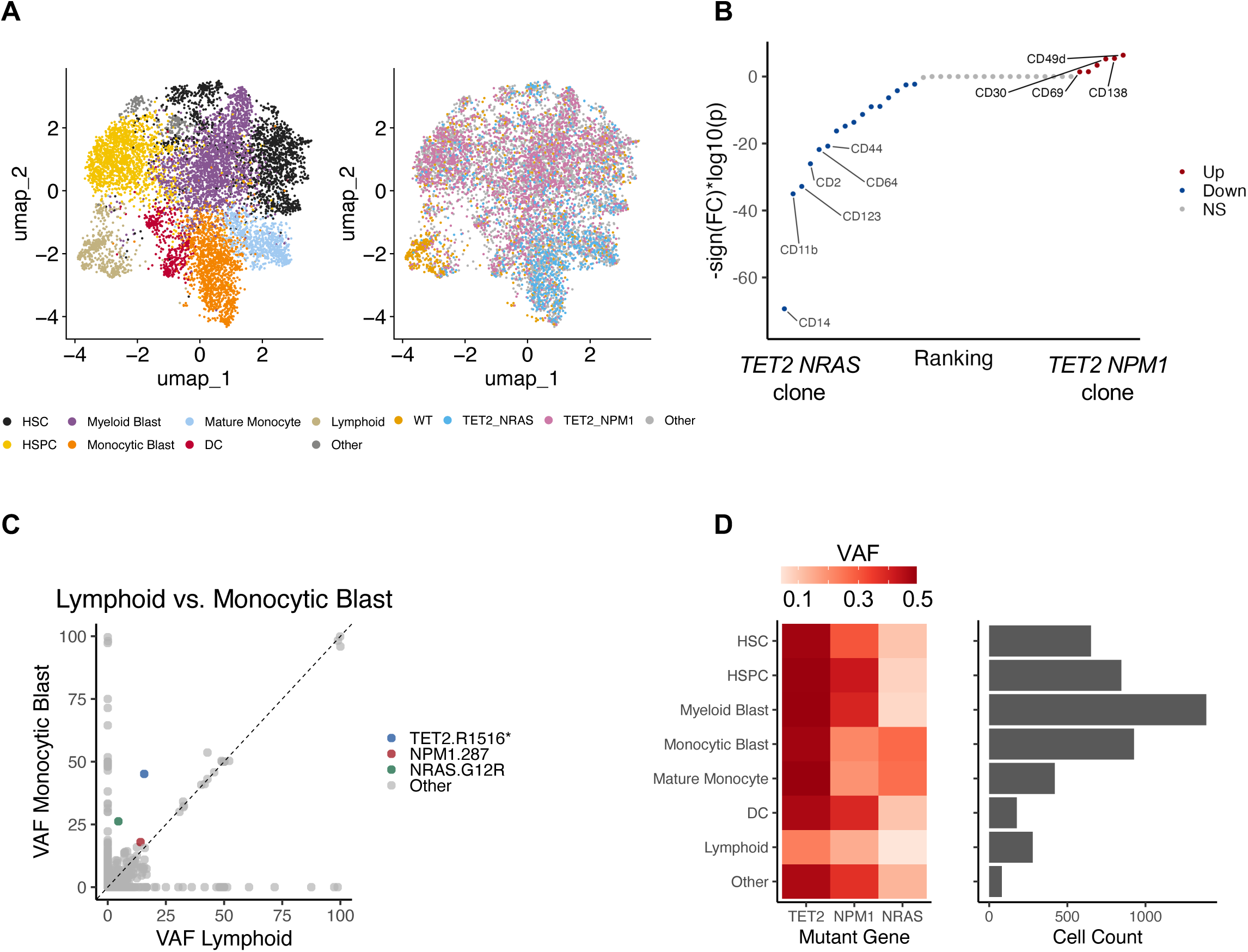
Identification of cell type-enriched mutations. **A)** UMAP of the cell types (left) for human acute myeloid leukemia sample harboring clones comprising 3 mutations (right: *NPM1*, *NRAS*, *TET2*). **B)** Dotplot depicting protein abundance difference between mutant clones. Wilcox significance (-log10(p)*sign of the fold change), is depicted on the y-axis, and proteins are ranked on differential enrichment between *TET2-NRAS* and *TET2-NPM1* on the x-axis (blue indicates upregulation toward *TET2 NRAS* clone, red indicates upregulation toward *TET2 NPM1* clone, and grey is neither). **C)** Scatterplot plot comparing VAF of lymphoid cells and monocytic blast population for each identified variant in the population. Above the dashed line, indicates a variant enrichment for monocytes while below would indicate variants enriched for lymphoid (*NPM1*: red, *NRAS*: green, *TET*2:blue). **D)** Heatmap depicting the VAF (red high, white low) for indicated mutations (x) for each cell type (y). Barplot depicts the cell count for each population in the sample.

Alternatively, a “cell type first” approach can use protein data to identify cell types (**Figure 3A**), grouping cell barcodes together before calculating variant allele frequencies for each cell population in bulk. In this example, we observed an enrichment of *NRAS* and *TET2* mutation abundance in in monocytic cells compared to lymphoid cells (**Figure 3C**). The enrichment plot can lead to downstream investigation of cell type specific variants, variants that are correlated, mutually inclusive, and mutually exclusive. This approach can also identify high VAF events that are restricted to rare cell populations, which might otherwise be mistaken for rare mutations or sequencing noise. Variants of interest, such as *TET2*, *NPM1*, and *NRAS*, can be compared across cell types to reveal larger patterns across the cohort of cells. In this example we observe monocytic cells have a higher VAF for *NRAS* compared to other cell populations (**Figure 3D**). In sum, these analysis approaches allow for direct inquiry of variants of interest across diverse cell populations or for discovery analyses of unknown variants by comparing cell types of interest.

### Active learning approach to sample demultiplexing

Sample multiplexing in single cell multi-omic sequencing is a powerful approach to reduce costs and batch effects. While there are many different demultiplexing strategies ^26–28^, we present a rapid strategy which builds off our prior doublet detection^29^, and does not rely upon external datasets from either antibody tagging or known variant information. Our demultiplexing approach is data-driven by first reducing the entire Allele Frequency (AF) matrix to full rank using an active learning approach to iteratively remove variants with low variance and cells with low genotyping quality. Next, k-means clustering is performed on this matrix to split the groups, with the cluster number defined by the user. An additional cluster is then added to account for cellular aggregates/doublets as in our prior doublet detection strategy^29^. Once clusters are defined, the cells that were excluded for low genotyping quality are imputed through a nearest neighbor approach to each cluster centroid. Cell clusters are then visualized by UMAP or heatmap for user-guided annotation with variants known to differ between samples. This allows for a data driven clustering of cells from each specimen and can be aided by sample-specific SNPs on panels, even if they are unknown to the investigator. Furthermore, this approach is robust to allele dropout of landmark SNPs, or underperformance of antibody derived tag (ADT) hashing.

We mixed four separate AML patient samples at different ratios to investigate whether the demultiplexing strategy could correctly identify disproportionately abundant clusters. We designed a panel that included amplicons targeting SNPs used for sample disambiguation 30. Each AML sample also contained unique combinations of somatic mutations including *STAG2, FLT3, IDH2, DNMT3A, NRAS, NPM1, KIT,* and *TET2.* Demultiplexing was then performed in two ways: 1) a baseline strategy consisting of a manually curated variant list based on SNP amplicons and expected variants for the mutated genes listed above, and 2) our proposed algorithmic variant discovery method (we denote as ML). We manually curated samples with exclusive somatic SNVs identified from bulk sequencing, including a mutation in *IDH2* only present in sample UP4 (**Figure 4AB**). We observed similar sample frequency between the two demultiplexing approaches (**Figure 4C)**, with the ML approach imputing a fraction of low-quality cells which were initially excluded from defining classes (1-18% for each class and a total of 10% of all cells). We observed a high sample assignment agreement between the two demultiplexing approaches (F1 scores: UP1=0.991, UP2=0.989, UP3=0.955, UP4=0.976, Doublet = 0.874) with significant deviation from random overlap (**Figure 4D**; Cohen’s κ = 0.965).

**Figure 4.**
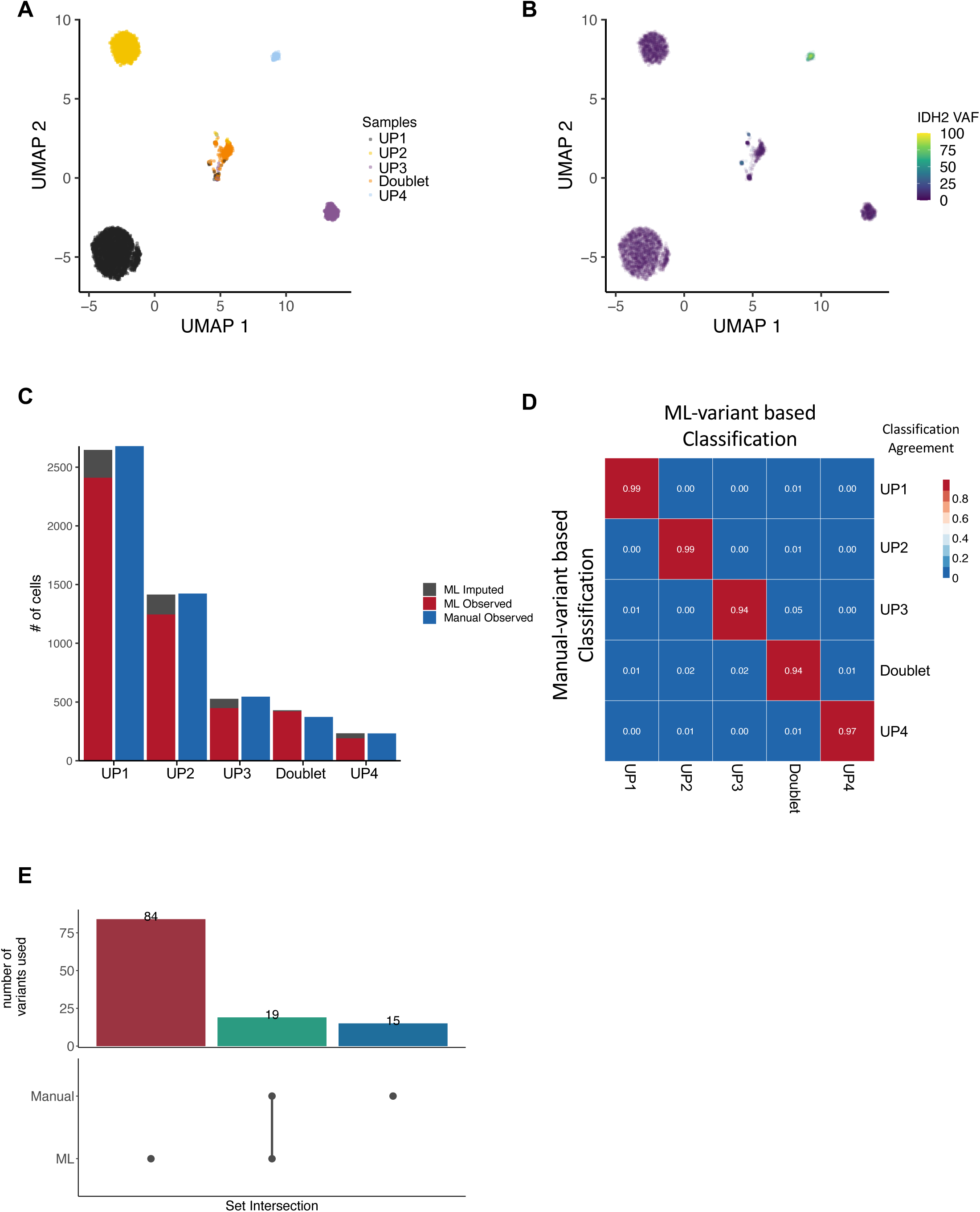
Active learning allows for informative feature selection and sample demultiplexing. **A)** UMAP depicting demultiplexing of individual samples following k-means clustering and manual label assignment. **B)** UMAP as in (A) with VAF for *IDH2* mutation overlaid showing enrichment in sample UP4. **C)** Barplot comparing the number of cells labeled with supervised strategies (blue), unsupervised machine learning (ML) strategy to determine the clusters (red), or poor-quality cells imputed by the ML strategy (grey). **D)** Confusion matrix for agreement of our supervised (manual) labeling and ML strategy containing a 4-sample multiplex. High agreement levels are along the diagonal. **E)** Upset plot for the overlap of variants used for each demultiplexing strategy.

We next assessed whether the unsupervised ML approach captures the same variants as our supervised list (**Figure 4E**). We identified 19 commonly identified variants between both methods, while the ML method uniquely found an additional 84 variants to aid in differentiating the samples. We found there was no significant overlap between the SNPs (hypergeometric p=0.187) and conclude that the ML approach removes the need for dedicated SNP amplicons for sample demultiplexing. Overall, the demultiplexing pipeline allows for a pooled samples to be split and downstream analysis to be performed.

### Copy Number Variation and Ploidy

The scDNA package has built in capabilities to determine copy number variations (CNV) along amplicons. We allow for flexible demarcation of a population of cells to serve as a diploid reference, a critical parameter for evaluating CNV in patient samples. We demonstrated this functionality with a *TP53-mutant* complex karyotype AML patient sample, where lymphoid cells were identified and noted as diploid reference cells. Using the methods described above, we identified cell types and *TP53* mutation status **(Figure 5A**). We observed lymphoid cells including T and NK were dominated by *TP53* WT cells, while myeloid cells harbored high *TP53-mutant* allele burden (VAF: 96.09%), reinforcing the choice of lymphoid cells as a CNV reference point. We next evaluated chromosome level ploidy across all cells, and identified a decrease in chr5 and gain in chr8 signal, matching the observed clinical karyotype of del5q;8+, common co-occurring genetic events in *TP53-*mutant AML (**Figure 5B**). We further refined these CNV observations across cell type (**Figure 5C**), observing significant differences in both cell type distribution of different chromosomal abnormalities (2-way ANOVA chromosome: F=554.786; p<0.001; cell type: F=5.787, p=1.01E-6). Post-hoc pairwise Wilcox-tests with a false discovery rate correction factor were then conducted for cell types per chromosome (**Supplemental Table 2**). We found lymphoid compartments were not altered for chr8 or chr5. As a representative example, HSPC-MPPs were statistically significant to NK cells in chr8(p=2.39E-20), chr5(p=1.04E-127), chr4(p=1.48E-10), and not statistically significant in chr13. We further find the only statistical significance between HSPC-MPP and HSCP-CDP are in chr4(p=2.85E-4). Collectively, these results demonstrate the capacity to identify cell type-specific chromosomal abnormalities.

**Figure 5.**
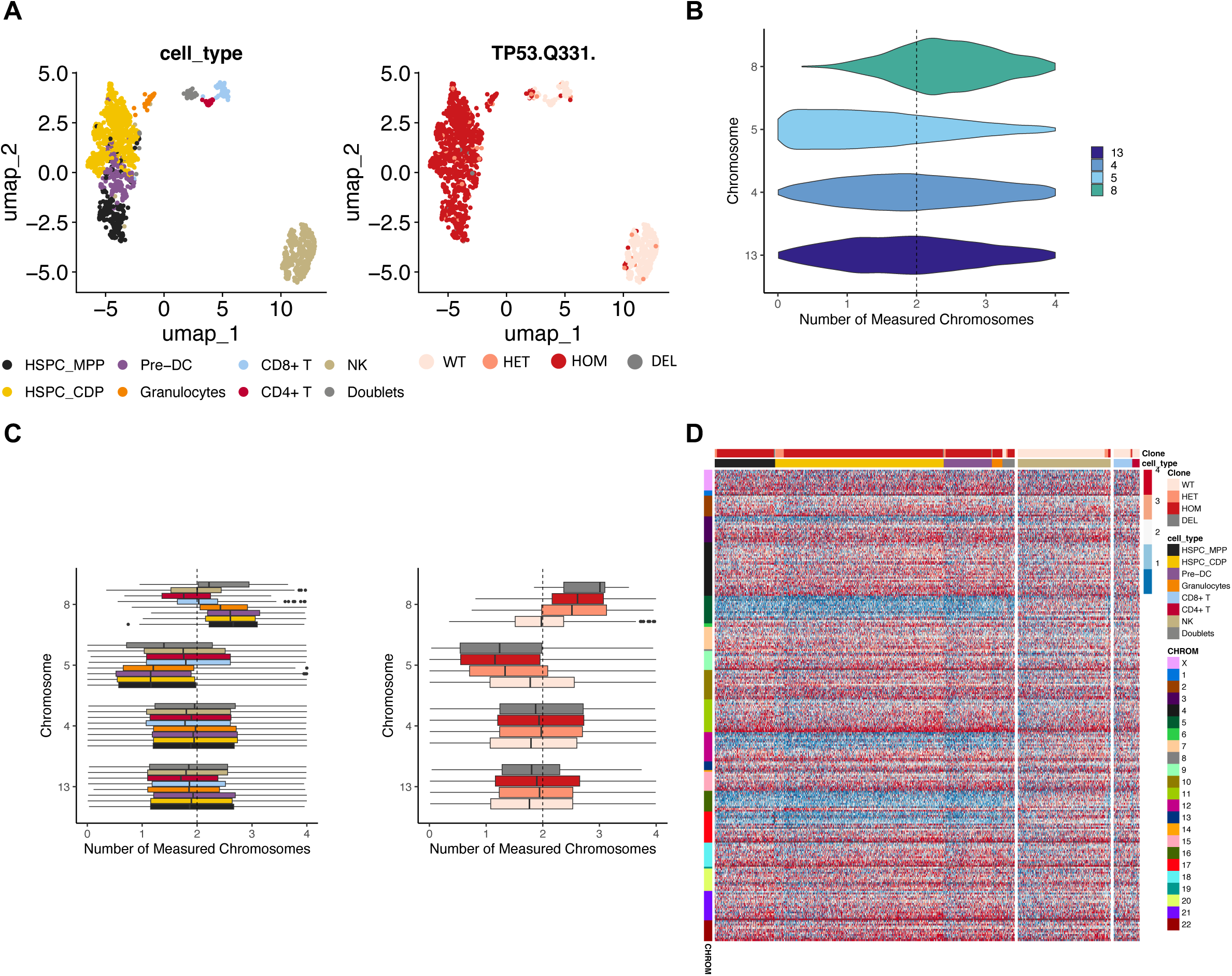
Inferring copy number variation associations with mutations and cell types. **A)** UMAP on underlying protein data showing cell types (left) and their *TP53* mutation status (right). **B)** Violin plot depicting chromosome abnormalities for the sample in (A) with a gain of chr8 and a loss of chr5. Chr4 and chr13 were chromosomes that have no anticipated alterations. **C)** Boxplots depicting CNV for cell types (left) and mutation status (right). **D)** Heatmap split between myeloid and lymphoid cell types for all amplicons. Individual cells are shown on the y-axis, and amplicons are arranged by chromosome on the y-axis. Blue indicates a loss of chromosomes, while dark red indicates a gain in chromosome.

We next compared CNV across *TP53-*mutation status and expectedly observed significant differences between WT and mutant cells (**Figure 5C**; F=4.483; p=3.76E-03; **Supplemental Table 2**). Wildtype cells did not possess chromosomal abnormalities; we found WT cells were significantly different from all other clones on chr5 and chr8 (p<0.001). Along chr13, the only significant difference was between WT and homozygous (p=9.00E-6). Critically, we also observed a subset of cells where *TP53-*mutant status could not be determined. We hypothesized that these cells might harbor deletion *TP53* on chr17 and evaluated CNV patterns across amplicons (**Figure 5D**). Within the myeloid cells, we observed a deletion of chromosome 17 where *TP53* mutation occurred. Furthermore, homozygous cells possessed decreased chr17 deletion, indicating a hemizygous state as opposed to a copy neutral loss of heterozygosity event. When considering CNV along cell types, we observed that most amplicons along chromosome 5 were lost in myeloid cells compared to lymphoid cells. Further abnormalities were found on chr12, and chr16 enriched in myeloid cells, indicating the lineage restriction of the complex karyotype AML sample. Overall, this approach enables investigation into SNV and CNV relationships in distinct cellular compartments, critical facets of understanding cancer cell evolution.

### Quantification of allele dropout events

The core output of our pipeline is represented in a plot inspired by oncoprints which we call a “clonograph.” Here we represent the constituent variants of a particular clone in a heatmap and the abundance of the clone in a bar plot (**Figure 6A**). In the example we build a clonograph for 3 mutations *TET2*, *NPM1*, and *NRAS,* where the most abundant clone is double mutant for heterozygous mutations in *TET2* and *NPM1*. We additionally observe rare clones which can arise as true genetic subclones, or a result of technical artifacts due to allele dropout (ADO) during PCR amplification. These events result in a true heterozygous cell being erroneously identified as wildtype (backward ADO) or homozygous (forward ADO). We assume that allele dropout is stochastic regarding the mutant allele and effects forward and backward events at the same rate.

**Figure 6.**
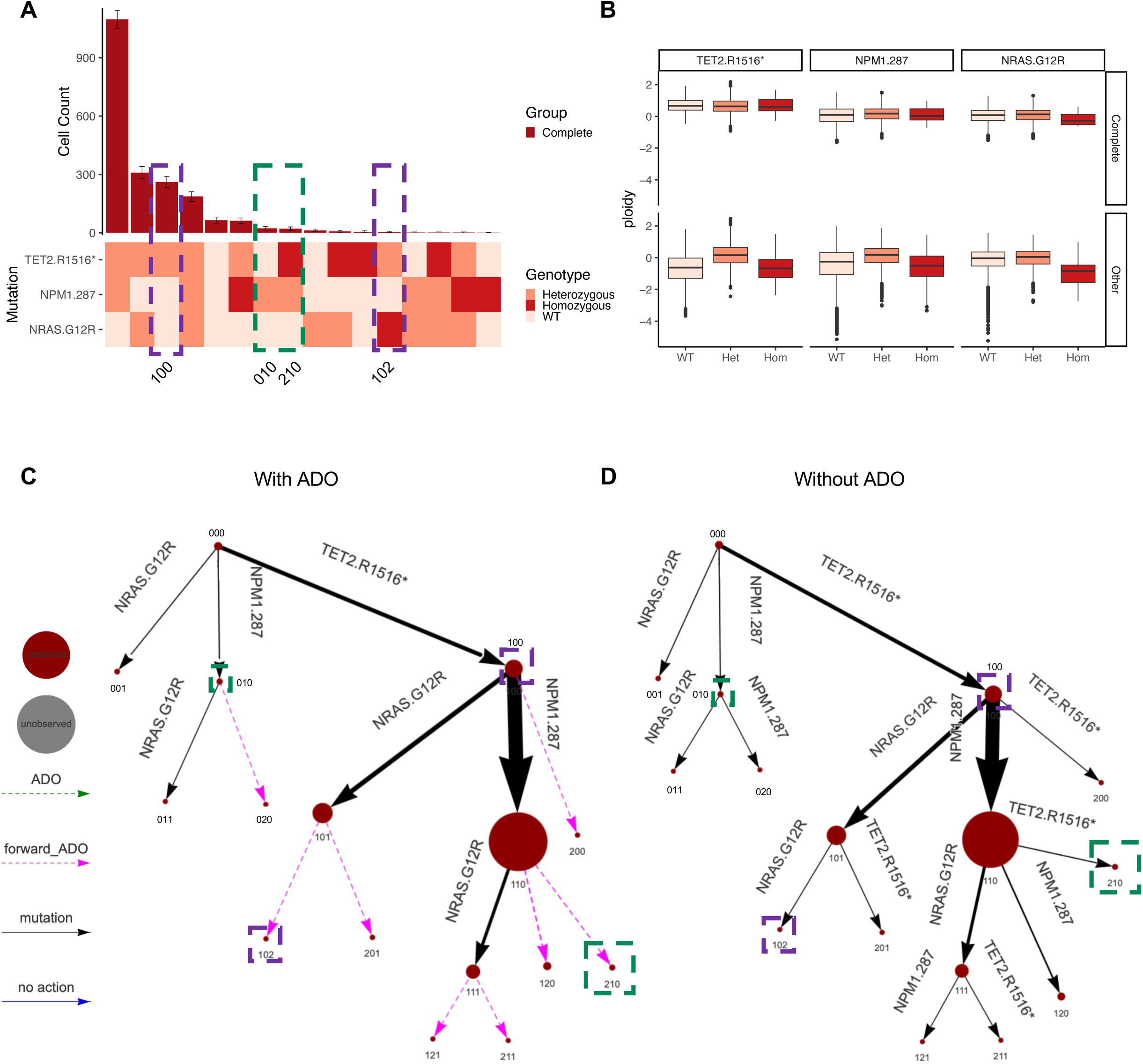
Reinforcement learning for an allele dropout aware trajectory analysis. **A)** Clonograph representing the clonal architecture of mutations in the heatmap, with bar graph indicating cell abundance for each clone. Light colors indicate WT, salmon is Heterozygous, and red is Homozygous. The purple and green boxes represent Allele Dropout (ADO) pairs. The green pairs are for the dominant clone, while the purple is for the second dominant clone. **B)** Boxplot of ploidy calculations for each variant split by “ passed QC calls (Complete) and cells that did not pass QC (Other). **C-D)** Clonal trajectories with (C) and without (D) consideration for ADO. The size of the dots are relative clone abundance, black lines indicate likely transition due to mutation, while pink lines are from forward ADO. Respective color boxes are around clones for ADO pairs in (A). Small clones are predominantly found to be results of ADO; however, some small clones are more likely to be a result of true mutations such as *NRAS* only, and *NPM1* and *NRAS* heterozygous clones. Subscripts indicate clone number in (A).

To demonstrate how ADO might influence clonal distribution, we highlight allele dropout pairs (both a forward and backward ADO event) with coded boxes in **Figure 6A**. The green boxes represent an ADO pair for the *TET2* mutation in the dominant clone (first bar). For this pair, nearly equal number of cells are found in both clones suggesting a similar ADO rate from a nearly ubiquitous *TET2* heterozygous mutation. In a second ADO pair, we turn to the 2nd most abundant clone harboring *NRAS* mutant cells indicated by purple boxes. In this setting, we observe a disproportionate bias towards the *NRAS^WT^* clone compared to the *NRAS^Hom^* clone. We hypothesize that the *NRAS^Hom^* clone is likely a result of ADO from *NRAS^Het^* cells, which manifest as a decrease in copy number. With the assumption that ADO directionality is stochastic, we further hypothesize that the *NRAS^WT^* clone is a mixture of true *NRAS^WT^* cells, and *NRAS^Het^* cells that underwent ADO and did not amplify the *NRAS^mutant^* allele. We rely on ploidy calculation and false positive rates to distinguish these possibilities (**Figure 6B**). We separated cells by quality classification, and observed uniform ploidy across genotype calls for *TET2* in high quality cells. In contrast, *NRAS^Hom^* and *NPM1^WT^* mutation calls were associated with a decrease in copy number, indicating a failure to amplify both mutant and WT alleles at each locus. Broader significant alterations were observed in low quality cells (**Supplemental Table 3**). To quantify the ADO toward either WT or homozygous calls, we calculated false positive rates using the ploidy calculations for all quality cells (see methods). We observed a false positive rate of 47.9% for *NRAS^WT^* and 86.8% for *NRAS^Hom^*. This overall tracks with expectations where a homozygous *NRAS* in AML is exceedingly rare.

### Mutation order through Reinforcement Learning

Finally, our pipeline provides tools for investigating the order of somatic mutation acquisition, also termed clonal trajectories. We use a Markov Decision Process (MDP) with clones treated as states and actions representing mutations and ADO events. We then perform Reinforcement Learning (RL) through Q-learning paradigms to identify the most likely trajectories to reach states. This approach is preset to identify ‘optimal’ clonal trajectories as we have previously reported but can also be altered to allow users to specify navigation to specific clones of interest such as the most dominant clone. We present this data as a clonal tree structure (**Figure 6C**). The green and purple boxes around circles correspond to their respective ADO pairs discussed above. We compared trajectories which use ADO false positive rates in the weighting against an ADO agnostic approach. Without considering ADO, all clones were interpreted as valid states, whereas with ADO, small clones that splinter from larger ones were assessed as unlikely states. These analyses reveal that in this sample, mutations in *TET2* likely precede mutations in *NPM1*. Including ADO in the model resolves that loss of heterozygosity of either *TET2* or *NPM1* does not exceed a false positive rate associated with copy number loss are likely artifacts of ADO. This output policy aligns with conventional models of clonal hematopoiesis associated-*TET2* mutations preceding leukemic transformation driven by *NPM1* and *NRAS*.

## DISCUSSION

We present a framework that is highly versatile for different applications of genotype immunophenotype data. The modularity of the SingleCellExperiment class used in the scDNA package offers critical flexibility to adapt with new emerging technologies, including single cell whole genome sequencing (scWGS)^31, 32^, scWGS+RNA^33, 34^, single cell targeted analysis of the methylome (scTAM-seq)^20^, scDNA+RNA^22, 23^ and scATAC+genotyping^35^. These emerging technologies can be readily adapted into our pipeline to identify clonal architecture and compute clonal statistics. Our pipeline is modular allowing for easy interfacing with new analytical approaches including normalization tools such as dsb, and visualization in Seurat^36, 37^. We will continue to integrate wrapper functions as new packages are developed to process proteogenomic data. We envision new iterations of scDNA will continue to push the frontier of single cell genotyping through additional investigation into mutation burden, subclonal diversity, and genotype-immunophenotype relationships. In its current form the package establishes a foundation to ensure quality assurance for a wide range of analyses.

This package adds to a growing suite of tools for scDNA analysis both in R and python^4-^ 6. This work has made significant advances in unifying variant annotation, genotype extraction, clone identification, and clonal trajectories. Furthermore, these different applications demonstrate how protein data can contribute to quality control, and immunophenotypic responses for analysis. Cell type specific VAFs, CNV, and clonal statistics can drive novel insights into cell type specific genomic evolution patterns not otherwise achievable through bulk analysis. The example applications shown are not theoretical examples of potential uses, as our open source package has already been deployed in a number of settings including: cohort analysis^7, 15, 24, 38^, rare event capture for clonal evolution and measurable residual disease detection^13, 39, 40^, and multi-omic studies^15, 41, 42^. While we present a series of vignettes to showcase a wide range of uses, not all examples are needed for a singular analysis. Toward this end, our vignettes are readily accessible on github (https://github.com/bowmanr/scDNA) to offer starting points for different types of analysis.

## Methods

The core structure of the multi-omics data is built upon the readily available SingleCellExperiment data object, shown in **Figure 1B**. The object allows for each cell barcode to have a container to hold summary statistics identified for each sequencing variant. The data includes layers for variant information including a Genotype calls (NGT) matrix, allele frequency, read depths, and genotyping quality. Additional information is embedded in the data object after completing different stages of analysis in our framework, including sample information, clone association, and cell confidence. Various metadata slots are used to store outputs of analysis. Within the analysis object, we contain droplet information for independent analysis, clonal architecture (a representation of the clones for the specific variants in the sample), clonal statistics, and trajectory analysis (e.g., the likely path mutations took from wildtype to reach clones of interest). Additionally, we store protein data and copy number variation data as AltExperiments within the SingleCellExperiment class where they can be interrogated with different normalization strategies. Producing the data layers to fill the SingleCellExperiment object is done by following our scDNA workflow as shown in **Figure 1A**. The overall workflow of the package is as follows:

1. Import data from Tapestri pipeline (H5) or a multi-sample VCF
2. Identify, annotate, and select variants of interest
3. Importing data into a single cell experiment object
4. Building clonal architecture
5. QC control including cell confidence labeling

Lastly, we deploy methods for demultiplexing pooled samples which recovers cells from each sample to be used independently in the pipeline. The following subsections explore the scDNA framework in more details. Examples of the pipeline are performed on hematopoiesis, specifically myeloid malignancies and acute myeloid leukemia (AML), with potential offshoot analysis that can be conducted using the scDNA package.

### Variant identification

The first phase of the scDNA package is to read in the NGT data (which contains four potential calls, wildtype, heterozygous, homozygous, missing) and identify the variants of interest that pass quality filters using a variant_ID() function. The filters are for *genotyping quality* (GT) and *variant allele frequency* (VAF). The genotyping quality cutoff is the sufficient and necessary number of cells that have successful calls for that gene. It does not matter whether the cells were called wildtype, heterozygous, or homozygous for this filter. It only compares whether an actual call or a Missing call was made. The second filter, based on VAF, determines whether the successful calls are mutated at a high enough capacity to warrant investigation.

Adjusting these two filters establishes an exploratory phase to find many variants of interest, or a refined search to narrow down the number of variants of interest. To ensure the exploratory phase is used, set the GT_cutoff=0, and VAF_cutoff=0. This will produce all possible variants that may be of interest, disregarding quality of the cells. The refinement phase is used after we have identified variants of interest and allows for quality control of cells harboring those variants. We recommend starting with an exploratory phase first before refinement phase (GT_cutoff=35 and VAF_cutoff=5).

The variants are annotated from nucleotide positions on the chromosomes to their standard gene nomenclature, then these are joined together into a single data frame. The current accepted annotation panels built into the package are the mm10, UCSC.hg19 and UCSC.hg38, as well as custom built TxDB panels. Additionally, we have a function that allows users to create their own TxDB with generate_txdb() function. It is encouraged to provide your own additional filtering to further refine the search such as class of mutations (Exonic, or Spicing, etc.) or more stringent refinements on the VAF. It is not strictly necessary to build a new TxDB for each panel, however it will decrease processing time.

### Extraction of variants to Single Cell Experiment object

The current accepted platforms for the data are h5 files from Tapestri and multiple sample VCF files. Information that is extracted from these files include Variant Allele Frequency (VAF), Allele Frequency bandpass (AF), Depth reading (DP), Genotyping Quality (GQ), and the NGT matrix. The AF filter is the allowable deviation from the WT, Het, or Hom call to be considered acceptable. For instance, assume we have an AF cutoff of 20 representing the total width of the band (so +/- 10%). Het is noted as a 50% for a particular gene, however, variants that fall with 40%-60% are treated as Het. DP is the minimum number of reads necessary for a reliable genotype call in a single cell. Sequential filtering refines a set of good quality cells for the variants of interest.

### Enumerating clones

After identifying the variants, the next step is to determine what clones exist and defining cell confidence labels to break down the quality of cells in each clone. To build the clones we devise a *clone code* based on the NGT matrix. A clone consists of genotype calls for the specific variants. We denote a clone as a singular column where each row index is a specific variant, and the value of the column (0=Wildtype, 1=Heterozygous, 2=Homozygous, 3=Missing) represents the genotype call. For instance, in the example shown in **Figure 6A**, the lighter to darker colors signal a WT to homozygous calls. We exclusively reject clones that contain Missing calls. The cells with the same clones are counted and sorted based on the abundance. The clonograph in **Figure 6A**, is assembled with the most dominant clone and decreasing order of abundance for the subclones.

Empirical confidence intervals are created through a bootstrapping strategy to enumerate clones. The bootstrapping approach resamples the entire clone, not a mix and match of the variant calls. Afterwards, a quantile range is established for the confidence interval. The bootstrapping is done for the cell confidence label based on cell quality. The first cell confidence labeling is based on passing quality filters from the data extraction that include the NGT (ensuring that no missing calls are made for the genes), from the AF filter (ensures allele calls are within suitable bounds), DP filter (sufficient read depth), and GQ filter (high enough genotyping quality). Passing all these filters refers to a “Complete” cell label. A cell is labeled as “Other” if the Genotype Analysis Toolkit (GATK) made a genotype call, but failing any one of these filters. **Figure 6A** only presents a clonograph with Complete cells.

### Multi-omics cell labeling

We investigate whether droplets are of high quality by comparing the overall DNA and protein content. Contour plots of **Figure 2A** shows most Empty droplets hold low DNA content, and lower protein content, however, there are lower quality candidates that overlap with the known droplets. Compared to empty droplets, candidate cells should have higher DNA and protein content. A more refined search in the Cell population allow for a more rigorous quality classification for each droplet. **Figure 2B** shows the outcome of our cell confidence classification. The classification has a hierarchical structure with three potential labels, poor DNA, low confidence, and high confidence. The first level is to look at DNA features to determine if a candidate cell has poor quality DNA. The second level incorporates protein information to determine the high or low confidence of a cell. Each droplet will be denoted as *c*, amplicon as *i*, protein as *j*, and the counts is represented as *x*.

### First level classification to determine poor DNA content

The classifier is built on a heuristic to evaluate the DNA content for each droplet. Two qualities are essential when considering cell’s DNA. The total counts of the amplicons, and the uniformity of those counts across the amplicons. A good quality cell with have high counts, and high uniformity compared to other droplets. To determine the uniformity, we first normalize the counts per amplicon to determine the compositional makeup for each droplet, *p*.(*x*_*c, i*_):

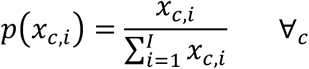

Using the normalized counts, the entropy of each cell, *H_c_*, is calculated:

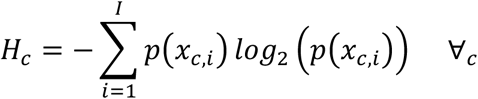

The efficiency measure is how we determine the uniformity, whereby a perfectly uniform distribution is used to determine the maximum theoretical entropy, *H_max_*. The efficiency/uniformity measure, *U_c_*, is as follows:

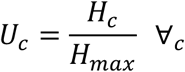

The uniformity measure of 1 means it is perfectly uniform, while a lower score indicates it is less uniform. Real cells are more likely to hold high uniformity. With these two measures the following heuristic is devised to score a cell for the DNA quality, *S_c_*.

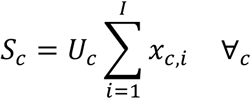

The score measure is high when cells have high counts that are similar across all amplicons. Higher scores indicate a higher chance of a droplet being classified as a cell, while low scores would indicate empty droplets. An empirical CDF is then produced for all droplets to extract upper tail p-values of the distributions, *p_s_c*. Droplets that are designated as a “Cell” from the Tapestri platform are investigated to determine if they are significantly above the bulk of the droplets by a threshold p-value, *p_thresh_*:

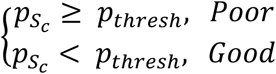

In the example **Figure 2B**, *p_thresh_* = 0.025. Since droplets are considered, not every sample will have a poor DNA quality. This is sample dependent and depending on the overlap between empty droplets and real cell populations. Once this level has been determined, the protein is considered on a second level.

### Second level classification to determine high and low confident cells

The primary purpose of the protein information in single cell sequencing is to distinguish cell types. The DNA assumptions for uniformity and total counts are ill equipped to handle the necessary requirements to quantify whether a cell has good or poor quality. Protein specific assumptions must be considered. For instance, in a 45-plex protein panel, we do not expect a cell to express all 45 markers. The implications that a droplet expresses all these markers is that it is a multi-cellular aggregate (doublet), or that a cell is dead, and permeabilized membranes have resulted in non-specific antibody uptake. On the other side, a low number of markers on a droplet requires context to determine whether it is noise or an actual cell. If a panel has heavy myeloid bias and relatively few lymphoid markers, the lymphoid population may appear as noise based on the markers since they should not hold myeloid markers. One key difference from DNA is that for proteins we do not expect panel uniformity, and total counts for each cell play a less critical role in determining their confidence. Towards the effort to classify them as high or low confidence, new measures are introduced to detect outliers within a cell.

The protein matrix is first turned into a pseudo count by adding a count of 1, and log-transformed to standardize the counts in a normal distribution rather than a skewed right distribution. This transformed protein is denoted as *y_c,j_*.

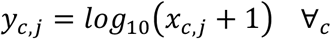

Next each cell is normalized based on the composition of the proteins to obtain probability measures for each protein.

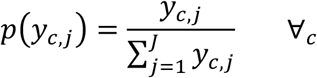

For each cell, the protein probability is sorted, and a cumulative distribution function is obtained through a cumulative summation. A reference CDF is built to model a perfectly uniform distribution. A uniform distribution is unlikely to be a real cell as this would imply that every protein marker is on the cell and often, markers are mutually exclusive to distinguish cell types. The L2 norm is calculated for each cell’s CDF and the reference distribution:

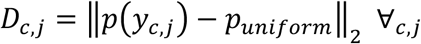

A second measure is to determine the smoothness of the CDF compared to the reference cell. This error term is the angle difference between the uniform distribution and each protein point in the cells’ CDF:

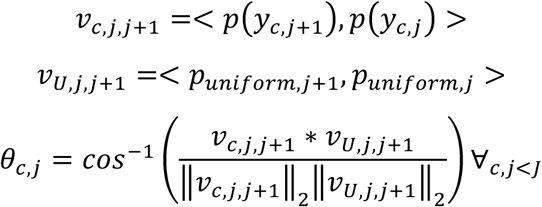

With these measures established, we follow an Empirical-Cumulative-distribution-based Outlier Detection (ECOD) algorithm^43^. First, we place all measures into a single matrix where the columns are the protein distance, protein smoothness, DNA quality heuristic, total protein counts, and rows are each cell.

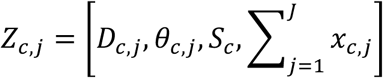

Next, we build both a left tail and right tail empirical CDF for each column which are denoted as 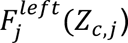, and 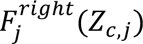. A skewness measure^44^ is used for each column to determine whether the cell population is left or right skewed, thus the appropriate empirical cdf can be applied.

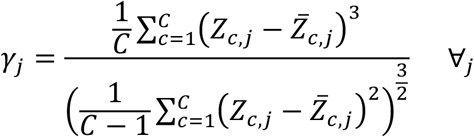

A positive skewness measure uses a right tail while a negative skewness uses a left tail. Each entry in the matrix has a probability value associated with it. The following equation is used to determine the outlier value for each component:

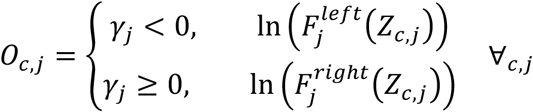

Lastly, a combination for a cell’s total outlier score is summed and is designed to find cells that exceed high levels of outlier score:

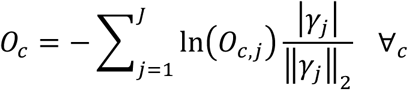

A cutoff value to determine inlier vs outlier cells is through a standard Tukey’s rule:

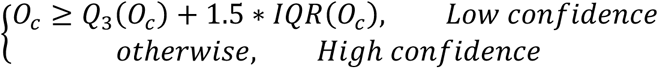

The cells with high levels of outlier scores labeled low confident cells, while those accepted are considered high confident. **Figure 2B** shows the outlier scores for different aggregated information of each cell, which includes DNA content, protein total, and protein markers expressed. The protein information demonstrate that higher markers expressed are more likely to be outliers. Likewise, cells that have lower marker expression also tend to have higher outlier scores. Cells that express all markers tend to carry more total protein content as well. These cells are labeled low confidence as they do not help in distinguishing cell types and induce noise in downstream clustering.

## Additional downstream analysis

### Demultiplexing

Our demultiplexing functionality starts by finding a reduced full rank Allele Frequency matrix through iterative thresholding. Poor quality cells and variants are discarded. The output matrix is then a reduced set of cells that contain variants which identify the different samples through a k-means clustering. The k is determined by the number of clusters expected and input by the user. We recommend adding an additional group to the k to capture a doublets population. Afterwards, the classification is imputed for the cells that were discarded by finding the shortest distance to the existing clusters. These cells are labeled for samples, split, and individually ran through the pipeline for genes of interest.

### Ploidy and CNV

One imperative analysis tool developed in this software package is using Copy Number Variations to assess ploidy, allele dropout and false positive rates of calls for specific genes. CNV is assessed from h5 files containing read depth information for each amplicon. First, we split cells into two groups, a *reference set* (*c* ∈ *R*) and *other set*. For each cell we then find the fraction (*f*_*i*,*c*_) of the read depth (*DP*_*i,c*_) overall all reads on that cell.

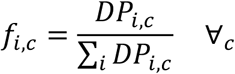

Then, the fractional values are standardized 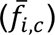 across amplicons through dividing by the median 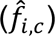.

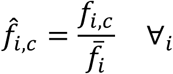

Ploidy is then calculated by the ratio of each cell’s standardized fraction 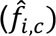 to the reference set cells 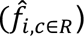 Afterwards we take the log2 of the of the ratio:

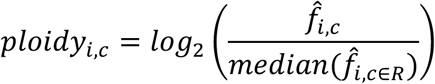

We use ploidy to produce standard t-test confidence intervals for heterozygous calls for each variant. The lower bound of the 99% confidence interval (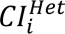, with a t-statistic of *t*_2.99_) shown below, where 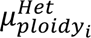 is the average ploidy for the heterozygous calls for the variant, 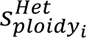 is the sample standard deviation, and *n^Het^* is the total heterozygous calls.

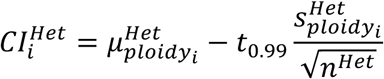

Afterwards, we count the number of wildtype (or homozygous) cells, *n^Het→Mut^*, that have a ploidy below the lower bound of the 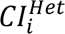.

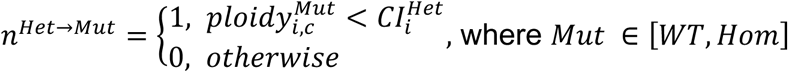

The false positive rate, α*^Het→Mut^*, is the ratio of cells that fall below the interval, *n^Het→Mut^*, over all cells for that mutation, *n^Mut^*,.

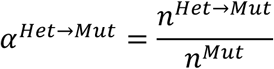

The false positive rates are then used as ADO risk for acquiring mutations which are treated as an alternative action for clonal trajectories.

### Clonal trajectory analysis

To find plausible trajectories clones develop, we employed the use of a Markov Decision Process (MDP) to determine all possible state transitions. We then use Reinforcement Learning (RL) to determine the likely paths subclones develop. Two available MDPs exist in our framework, one which does not consider ADO, and another that accounts for ADO by adding an additional forward and backward path for heterozygous clones. We have prebuilt trajectories that are commonly used for analysis. These include finding 1) all optimal paths from the WT clone to every clone in the MDP, 2) the optimal tree to connect all observed clones as seen in clonograph, 3) all optimal paths from every observed clone to the most dominant clone, and 4) the most efficient path to the dominant clone from the WT clone. We also provide visualization of mutation order in a bubble plot as previously developed^45^.

The MDP is defined as a tuple {*S, A, R, γ*}. States (*s* ∈ *S*) are represented by the theoretical clones. Actions (*a* ∈ *A*) represent mutations or ADO events from one clone to another. *R* is the reward matrix for transitioning from state to state through an action. *R*(*s*^J^|*s*, *a*) is the reward function to give the reward after the transition from state *s* to state *s*′ with action *a* and discount factor γ to scale the future rewards to ensure boundedness. *R* is determined by clonal composition of a sample. Reaching an observed clone state is rewarded based on the reached states compositional abundance (*s*′*_CA_*). Rewards for ADO events are different depending on whether it is heterozygous to wildtype, or heterozygous to homozygous. The heterozygous to wildtype ADO event (*a* = *ADO^Het→Mut^*), appears as a “loss of mutation,” which is negatively rewarded to encourage trajectories to gain mutations rather than losing mutations. Conversely, heterozygous to homozygous ADO events (*a* = *ADO^Het→Hom^*) are positive rewards as they appear as a “gain in a mutation,” however, the reward is reduced compared to gain of a mutation at an alternative genomic position. Both ADO events are influenced by the false positive rates, α*^Het→WT^* and α*^Het→Hom^*, discussed in the CNV and ploidy section. The reward design is as follows:

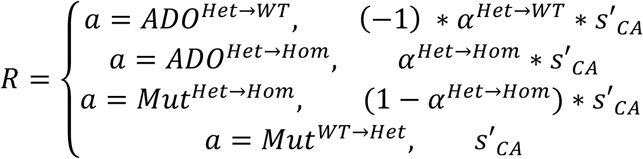

The employed RL strategy is based on Q-learning, a model-free temporal differencing approach, which relies on data, sufficient sampling, and random actions to update the policies and weighting schemes. The Q value is updated as follows:

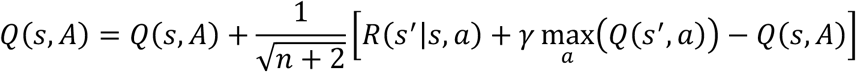

Where the *n*, is the sampling trajectory, and the entire term is the learning rate. The result is a converged Q-matrix which provides optimal policies for the entire MDP network. This Q-matrix is then used to find shortest paths, or desired paths for optimal order of mutations.

## Acknowledgements

R.L.B. is supported by a National Cancer Institute grant (R00 CA248460), an ASH Junior Faculty Scholar award and the V Foundation. L.A.M. is supported by a National Cancer Institute grant (R00 CA252005) and an American Society of Hematology (ASH) Junior Faculty Scholar award.

## Author Contributions

R.L.B. and M.B. conceptualized studies, designed and optimized bioinformatic workflow. L.A.M., S.F.C., O.S, and S.I.G designed single-cell DNA/Protein experimental methodologies. M.B., R.B., T.R., and R.L.B. performed all computational multiomic analysis. M.B., V.S, A.Z., S.G., and R.L.B generated manuscript figures and performed statistical analysis. L.A.M., O.S, S.I.G, S.F.C, and R.L.L provided critical data for analysis. M.B. and R.L.B wrote and edited the manuscript with edits contributed from L.A.M. All authors read the manuscript and agreed on the final version.

## Competing Interests

L.A.M. and R.L.B. had previously received honoraria for speaking arrangements and had previously served on a Speakers Bureau for Mission Bio, Inc. S.I.G has stock ownership interests in Carisma Therapeutics; advisory role with Asher Bio; research funding from Carisma Therapeutics and Novartis; holds patents for chimeric antigen receptor T cells for actue myeloid leukemia. S.F.C. has been a consultant and/or shareholder for Daiichi-Sankyo and Ursamin, none of which are directly related to the content of this paper. S.F.C. has received research support from Syndax, Trio, and Actinium. S.F.C. is an inventor on a patent related to Menin inhibition WO/2017/132398A1. R.L.L is on the Supervisory board of Qiagen (compensation/equity), a co-founder/board member at Ajax (equity), a board member of the Mark Foundation for Cancer Research and is a scientific advisor to Mission Bio, Kurome, Syndax, Scorpion, Zentalis, Jubilant, Auron, Prelude, and C4 Therapeutics; for each of these entities he receives equity/compensation. He has received research support from the Cure Breast Cancer Foundation (with IP rights), Calico, Zentalis and Ajax, and has consulted/provided professional services for ECOG-ACRIN, Genome Canada, Goldman Sachs and Astra Zeneca.

## Data Sharing Statement

Raw data files are available upon request to the corresponding authors and will be made available on dbGAP prior to final publication.

## Code Availability

All scripts and software are publicly available at https://github.com/bowmanr/scDNA/.

---

**Supplemental Table 1: QC metrics available.**

**Supplemental Table 2: CNV P-values for amplicons and cell types for ploidy.**

**Supplemental Table 3: ADO boxplot statistical analysis and False positive rates**

## Notes

https://github.com/bowmanr/scDNA

## References

1. Heumos, L. et al. Best practices for single-cell analysis across modalities. Nat Rev Genet 24, 550–572 (2023).

2. Stuart, T. et al. Comprehensive Integration of Single-Cell Data. Cell 177, 1888–1902 e1821 (2019).

3. Wolf, F.A., Angerer, P. & Theis, F.J. SCANPY: large-scale single-cell gene expression data analysis. Genome Biol 19, 15 (2018).

4. Pei, D., Griffard, R., Yellapu, N.K., Nissen, E. & Koestler, D.C. optima: an open-source R package for the Tapestri platform for integrative single cell multiomics data analysis. Bioinforma2cs 39 (2023).

5. Wind, S.M., Reinkens, T., Behrens, Y.L. & Sandmann, S. scafari: exploring scDNA-seq data. Bioinforma2cs 41 (2025).

6. Mays, J.C., et al. KaryoTap Enables Aneuploidy Detection in Thousands of Single Human Cells. bioRxiv (2024).

7. Miles, L.A. et al. Single-cell mutation analysis of clonal evolution in myeloid malignancies. Nature 587, 477–482 (2020).

8. Morita, K. et al. Clonal evolution of acute myeloid leukemia revealed by high-throughput single-cell genomics. Nat Commun 11, 5327 (2020).

9. Schwede, M. et al. Mutation order in acute myeloid leukemia identifies uncommon paderns of evolution and illuminates phenotypic heterogeneity. Leukemia 38, 1501–1510 (2024).

10. Poon, G. et al. Single-cell DNA sequencing reveals pervasive positive selection throughout preleukemic evolution. Cell Genom 5, 100744 (2025).

11. Stiff, A., et al. Multiomic profiling identifies predictors of survival in African American patients with acute myeloid leukemia. Nat Genet 56, 2434–2446 (2024).

12. Iacobucci, I. et al. Modeling and targeting of erythroleukemia by hematopoietic genome editing. Blood 137, 1628–1640 (2021).

13. Robinson, T.M. et al. Single-cell genotypic and phenotypic analysis of measurable residual disease in acute myeloid leukemia. Sci Adv 9, eadg0488 (2023).

14. Ediriwickrema, A. et al. Single-cell mutational profiling enhances the clinical evaluation of AML MRD. Blood Adv 4, 943–952 (2020).

15. Drucker, M. et al. Genotype-immunophenotype relationships in NPM1-mutant AML clonal evolution uncovered by single cell multiomic analysis. bioRxiv (2024).

16. Peretz, C.A.C. et al. Single-cell DNA sequencing reveals complex mechanisms of resistance to quizartinib. Blood Adv 5, 1437–1441 (2021).

17. Amezquita, R.A. et al. Orchestrating single-cell analysis with Bioconductor. Nature Methods 17, 137–145 (2020).

18. Demaree, B. et al. Joint profiling of DNA and proteins in single cells to dissect genotype-phenotype associations in leukemia. Nat Commun 12, 1583 (2021).

19. Ruff, D.W., Dhingra, D.M., Thompson, K., Marin, J.A. & Ooi, A.T. High-Throughput Multimodal Single-Cell Targeted DNA and Surface Protein Analysis Using the Mission Bio Tapestri Platform. Methods Mol Biol 2386, 171–188 (2022).

20. Bianchi, A. et al. scTAM-seq enables targeted high-confidence analysis of DNA methylation in single cells. Genome Biol 23, 229 (2022).

21. Scherer, M. et al. Clonal tracing with somatic epimutations reveals dynamics of blood ageing. Nature 643, 478–487 (2025).

22. Lindenhofer, D. et al. Functional phenotyping of genomic variants using joint multiomic single-cell DNA-RNA sequencing. Nat Methods 22, 2032–2041 (2025).

23. Yuan, D.J. et al. Genotype-to-phenotype mapping of somatic clonal mosaicism via single-cell co-capture of DNA mutations and mRNA transcripts. bioRxiv (2024).

24. Lee, D., et al. Prognostic and Therapeutic Implications of BRAF Mutations in Acute Myeloid Leukemia. bioRxiv (2025).

25. Van der Auwera, G.A. & O’Connor, B.D. Genomics in the cloud: using Docker, GATK, and WDL in Terra. (O’Reilly Media, 2020).

26. Kennedy, V.E., et al. SNACS: a tool for demultiplexing single-cell DNA sequencing data. Bioinforma2cs 41 (2025).

27. Borgsmüller, N., et al. *DemoTape*: Computational demultiplexing of targeted single-cell sequencing data. bioRxiv, 2024.2012.2006.627152 (2024).

28. Sollier, E., Kuipers, J., Takahashi, K., Beerenwinkel, N. & Jahn, K. COMPASS: joint copy number and mutation phylogeny reconstruction from amplicon single-cell sequencing data. Nat Commun 14, 4921 (2023).

29. Robinson, T.M. et al. Single-cell genotypic and phenotypic analysis of measurable residual disease in acute myeloid leukemia. Science Advances 9, eadg0488 (2023).

30. Pakstis, A.J. et al. SNPs for a universal individual identification panel. Hum Genet 127, 315–324 (2010).

31. Yin, Y. et al. High-Throughput Single-Cell Sequencing with Linear Amplification. Mol Cell 76, 676–690 e610 (2019).

32. Gonzalez-Pena, V. et al. Accurate genomic variant detection in single cells with primary template-directed amplification. Proc Natl Acad Sci U S A 118 (2021).

33. Olsen, T.R. et al. Scalable co-sequencing of RNA and DNA from individual nuclei. Nat Methods 22, 477–487 (2025).

34. Natu, A., et al. Single cell whole genome and transcriptome sequencing links somatic mutations to cell identity and ancestry. bioRxiv (2025).

35. Izzo, F. et al. Mapping genotypes to chromatin accessibility profiles in single cells. Nature 629, 1149–1157 (2024).

36. Stuart, T. et al. Comprehensive Integration of Single-Cell Data. Cell 177, 1888–1902.e1821 (2019).

37. Mulè, M.P., Martins, A.J. & Tsang, J.S. Normalizing and denoising protein expression data from droplet-based single cell profiling. Nature Communica2ons 13, 2099 (2022).

38. Fobare, S., et al. PTPN11 Mutation Clonal Hierarchy in Acute Myeloid Leukemia. bioRxiv (2024).

39. Stahl, M. et al. Molecular predictors of immunophenotypic measurable residual disease clearance in acute myeloid leukemia. Am J Hematol 98, 79–89 (2023).

40. Xiao, W. et al. A JAK2/IDH1-mutant MPN clone unmasked by ivosidenib in an AML patient without antecedent MPN. Blood Adv 4, 6034–6038 (2020).

41. Bowman, R.L. et al. In vivo models of subclonal oncogenesis and dependency in hematopoietic malignancy. Cancer Cell 42, 1955–1969 e1957 (2024).

42. Sande, C.M. et al. ATM-dependent DNA damage response constrains cell growth and drives clonal hematopoiesis in telomere biology disorders. J Clin Invest 135 (2025).

43. Li, Z. et al. Ecod: Unsupervised outlier detection using empirical cumulative distribution functions. IEEE Transac2ons on Knowledge and Data Engineering 35, 12181–12193 (2022).

44. Hornik;, D.M.E.D.K. & Leisch, A.W.F. (2024).

45. Malikic, S., Jahn, K., Kuipers, J., Sahinalp, S.C. & Beerenwinkel, N. Integrative inference of subclonal tumour evolution from single-cell and bulk sequencing data. Nature Communica2ons 10, 2750 (2019).

